# A Method for Computationally Constructing Eukaryotic Synthetic Signal Peptide Sequences

**DOI:** 10.1101/2021.11.19.469281

**Authors:** Grant T. Daly, Aishwarya Prakash, Ryan G. Benton, Tom Johnsten

## Abstract

We developed a computational method for constructing synthetic signal peptides from a base set of signal peptides (SPs) and non-SP sequences. A large number of structured “building blocks”, represented as *m*-step ordered pairs of amino acids, are extracted from the base. Using a straightforward procedure, the building blocks enable the construction of a diverse set of synthetic SPs that could be utilized for industrial and therapeutic purposes. We have validated the proposed methodology using existing sequence prediction platforms such as Signal-BLAST and MULocDeep. In one experiment, 9,555 protein sequences were generated from a large randomly selected set of “building blocks”. Signal-BLAST identified 8,444 (88%) of the sequences as signal peptides. In addition, the Signal-BLAST tool predicted that the generated synthetic sequences belonged to 854 distinct eukaryotic organisms. Here, we provide detailed descriptions and results from various experiments illustrating the potential usefulness of the methodology in generating signal peptide protein sequences.

---

Signal peptides (SPs) are protein sequences that play a role in protein secretion, transport, and trafficking within the cell^1^. SPs, sometimes referred to as a localization or targeting sequences, are short chains of 10 – 70 amino acids (AAs) that are typically present at the N-terminus of newly synthesized proteins and participate in the cellular localization or transport of the protein. Over the past twenty years, numerous software tools have been developed to identify such peptide sequences, but only recently have researchers started to develop computational methods to generate synthetic signal peptides (SSPs)^2–5^. The ability to generate novel signal peptides could advance synthetic protein development for industrial and pharmaceutical applications. An example of the utility of generating SSPs to target proteins to specific cellular organelles was observed with the successful targeting of an antioxidant protein to the mitochondria to alleviate oxidative stress within the organelle that otherwise leads to the development of neurodegenerative disorders such as Parkinson’s Disease^6^.

The vast majority of existing methods designed to generate SSPs are based on a neural network learning algorithm^2–5^. Limitations of these current algorithms include opaqueness in the generation of the SSPs (*i.e.* not knowing how or why a particular SSP was generated) and the requirement for large training datasets. In response, we have developed a computational method that is able to construct SSPs from small sets of known base SP and non-SP sequences. The method is significant in three ways. First, the building blocks (BBs) used to construct sequences are simply a combination of the twenty canonical AAs and as such are easily recognized by biologists and other scientists. Second, there is a high probability that a constructed sequence is classified as an SP by existing prediction platforms (Signal-BLAST, MULocDeep)^7, 8^. Third, the process facilitates the construction of large numbers of SSPs that collectively have a measurable degree of similarity with a wide range of organisms.

## RESULTS and DISCUSSION

The current method is based on the previously developed Multi-Layer Vector Space (MLVS) model^9^. It consists of three main steps: 1) discovery of candidate BBs, 2) selection of candidate BBs satisfying a threshold requirement (*i.e.,* qualified BBs), and 3) assembly of qualified BBs to create new SSPs (Fig. 1). These steps are performed in the context of a base set of SP and non-SP sequences. First, we identify *m*-step ordered pairs of AAs that occur at least one time in a base sequence, where *m* represents the number of spaces between AAs. All the ordered pairs made up of consecutive AAs form the 1-step ordered pair, *P*_1_. A MLVS is the set of *m*-step ordered pairs, *P*_1_,*P*_2_,…,*P*_k_. For instance, the *ASLGV* sequence contains the 3-step ordered pair [*S*, *V*] where *S* represents the anchor AA occurring at location 2 and *V* represents the tail AA occurring 3 locations to the right of *S*. Informally, the number of *m*-step ordered pairs of AAs that can be derived from a set of protein sequences is a function of the length of the sequences and number of distinct AAs contained within those sequences.

**Fig. 1:**
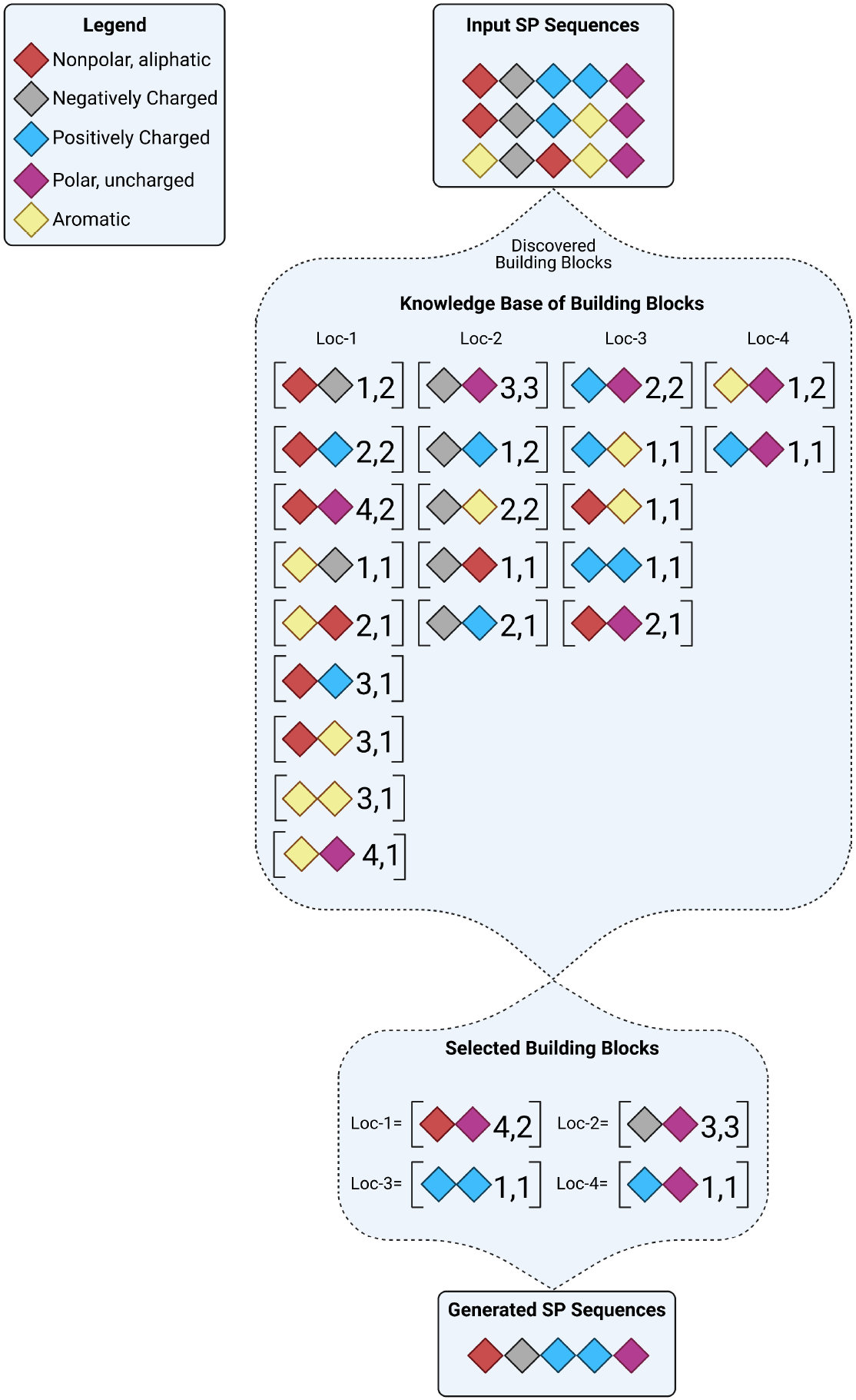
Overview of SSP generation. The process takes a set of SP and non-SP base sequences and constructs a knowledge base of BBs. The BBs are defined in terms of *m*-step ordered pairs of AAs, where *m* represents the number of spaces between AAs. Within each bracket two AAs are represented, followed by two numbers where the first number represents the location (loc) of the second AA from the first AA, while the second number represents the frequency with which this pair occurs. Colored diamonds represent the five broad classifications of AAs. BBs are discovered for each loc across the length of the base SPs. Novel SSPs are generated by selecting a BB for each loc. Created with BioRender.com.

Next, we identify a subset of candidate BBs used to construct new SSPs. We filter candidate BBs based on the absolute difference in the normalized frequency of occurrence of an *m*-step ordered pair across the SP and non-SP base sequences. The normalized frequency of a BB with respect to non-SP and SP sequences is equal to its frequency of occurrence divided by the number of non-SP and SP sequences, respectively. Those candidate BBs having an absolute difference value greater than a user-defined threshold between 0 and 1 are selected. The resulting set of qualified BBs are those that occur frequently in the SP base sequences and infrequently in the non-SP sequences, and vice-versa, allowing for increased diversity. This approach is inspired by the data mining concept of contrast patterns^10^. For example, assume we have a base set of 10 SP sequences and a base set of 12 non-SP sequences; further assume the user specified BB difference threshold is 0.6. If the 3-step ordered pair [S,V] occurs 7 times in the SP base set and 1 time in the non-SP base set, it then has a normalized frequency of 0.7 and 0.083, respectively. The absolute difference is 0.617, which is higher than the given threshold (0.6) and hence the 3-step ordered pair [S,V] would represent a qualified BB.

Finally, we use the qualified BBs to generate new SSPs. In constructing an SSP sequence *S* of length *n*, denoted as (*s*_1_, *s*_2_,…, *s*_n_), each location *s*_i_ (1 ≤ *i* ≤ *n*) is assigned all qualified *m*-step ordered pairs of AAs where the upper bound on the step size *m* is (*n* - *i*). The qualified BBs at each location *s*_i_ are sorted in non-decreasing order based on their frequency of occurrence across the base SPs. The frequency of a BB with a step size of *m* at location *s*_i_ is the number of SP base sequences that contain the BB anchored at location *s*_i_ (Fig. 1). Assembling a new SSP starts with the selection of a BB at sequence location 1, followed by location 2, and terminating at location (*n* – 1). BBs are selected at each location based on an input parameter, called the rank-position range, which represents a range of integer numbers from a specified lower to an upper bound. For each sequence location *i*, a value *v* within the given rank-position range is randomly chosen and the BB listed at position *v* is selected. Once a BB of step size *m* is selected at location *i*, its inclusion in the SSP is determined by the following conditions: If an AA already exists at location *i* and it matches the anchor AA of the selected BB, then the tail AA is assigned to the location (*i* + *m*) if it is currently unoccupied. If the location *i* is unoccupied, then the anchor AA is assigned to location *i*; in this case, the tail AA is also inserted at location (*i* + *m*) if the tail location is currently unoccupied. In all other cases, the anchor and/or tail AAs of the BB are not inserted into the evolving sequence.

We conducted a set of experiments to determine the accuracy and diversity provided by our method in generating SSPs (Fig. 2). Candidate BBs were discovered from 2,311 SP and 7,384 non-SP eukaryotic base sequences. The MULocDeep prediction tool identified 98% of the base SP sequences as being secreted to the extracellular space ^8^. A total of 27,213 qualifying BBs were generated based on an absolute difference value greater than zero. We generated different sets of sequences from these BBs using rank-position ranges [1,1000], [1001,2000], [2001,3000], and [3001,4000]. For each range, 500 sequences of 70 AAs were generated. For each range, we display the percentage of generated sequences defined by Signal-BLAST as SSPs (Fig. 2a) over 5 separate experiments. The MULocDeep tool identified 98% of the SSPs also as being secreted to the extracellular space^8^. The accuracy (*i.e.*, likelihood of generating an SSP) of higher-ranked positions is greater because the BBs are more likely to be found in the base SPs. We conclude from the experimental results that accuracy is dependent on the rank-position. We also calculated the accuracy of a naive method for generating SSPs that uses single AAs as BBs (Fig. 2a). The overall accuracy of this alternative method is much lower and the number of SSPs that can be generated is quite small in comparison.

**Fig. 2:**
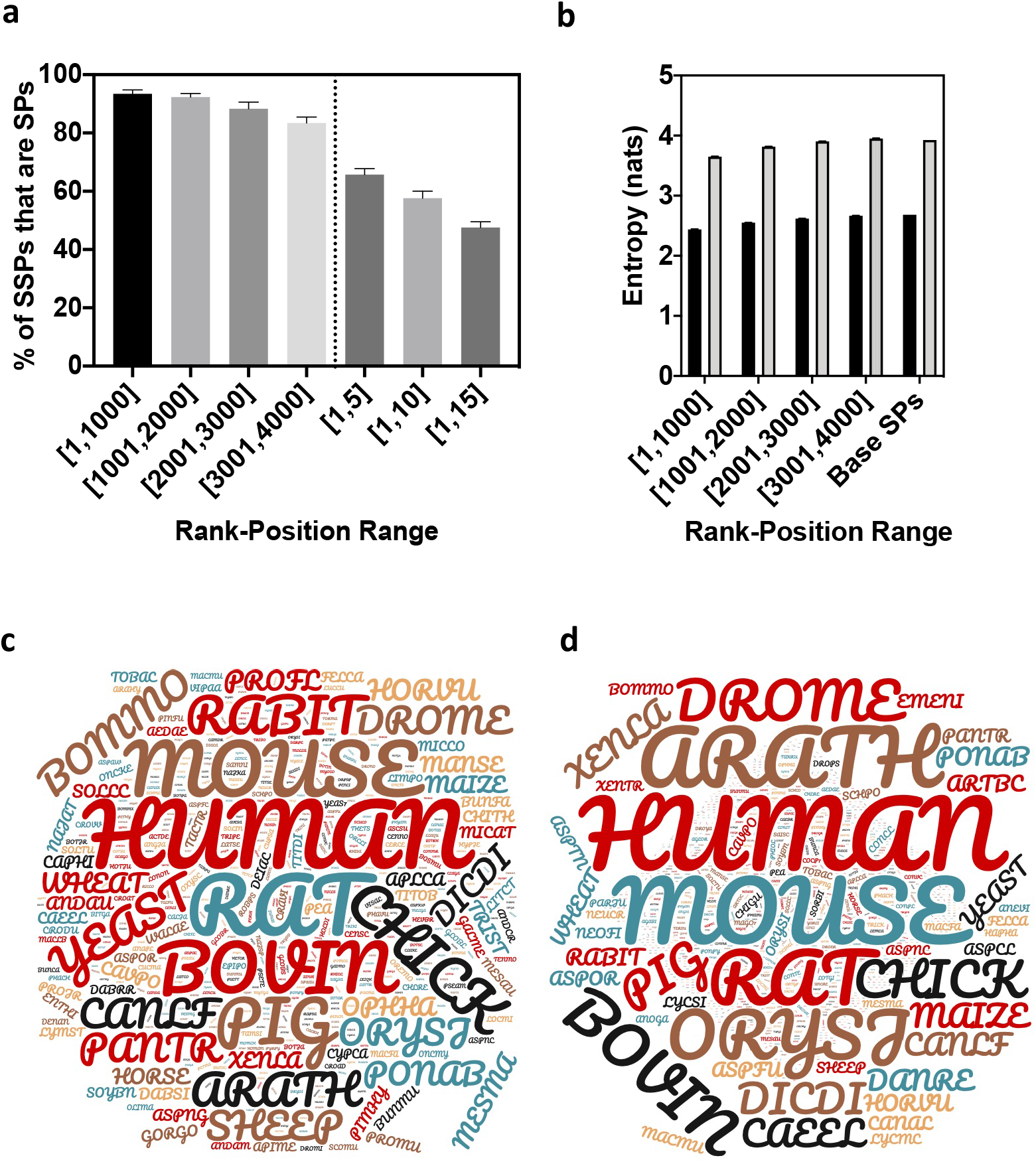
Validation of obtained SSPs. **a.** Accuracy assessment (5 replicates) for generated SSPs by rank-position range that were deemed SPs by Signal-BLAST. Accuracy results for the proposed method are given for ranges [1,1000], [1001,2000], [2001,3000], and [3001,4000] and accuracy results for the naive method are given for ranges [1,5], [1,10], and [1,15]. **b.** Entropy of single AAs (black) and bigrams (grey), represent the complexity of the base data and the generated SSPs. **c-d.** Organism diversity of the base SPs (**c**) and the generated SSPs (**d**) created using wordclouds.com. All organisms are displayed by shrinking the word cloud such that no words were omitted.

Next, we evaluated the entropy for the SSPs generated by our proposed method in each rank-position using the standard Shannon entropy measure for unigrams and bigrams. SSP were plotted as the mean entropy in “nats” for the 5 runs displayed as mean ± standard deviation (Fig. 2b), with values for base sequences included for comparison. Each rank position displayed low standard deviation, demonstrating batch consistency. Fig. 2b demonstrates that entropy increases as the rank-position value increases for unigrams and bigrams, with the [3001,4000] set having similar entropy to the base sequences. We concluded that the randomness of sequences is a function of the rank-position parameter, and that a user could “tune” this parameter to optimize accuracy at the expense of randomness and novelty. Extended entropy statistics are provided in Table 1. Lastly, we generated 9,555 sequences using a rank-position range of [1,3000]. According to Signal-BLAST, a total of 8,444 (88.37%) are defined as SSPs and covered a diverse range of organisms similar to that of the base SPs (Fig. 2c-d).

**Table 1:** Entropy Statistics for SSPs

| Entropy Statistics |  |  |  |  |
| --- | --- | --- | --- | --- |
|  | # Residues | # Runs | Mean (nats) | std |
| **1-1000** | **1** | 5 | 2.439357 | 0.004858 |
|  | **2** | 5 | 3.649838 | 0.005875 |
| **1001-2000** | **1** | 5 | 2.550365 | 0.003772 |
|  | **2** | 5 | 3.816099 | 0.003658 |
| **2001-3000** | **1** | 5 | 2.617792 | 0.003151 |
|  | **2** | 5 | 3.903525 | 0.003077 |
| **3001-4000** | **1** | 5 | 2.665958 | 0.003725 |
|  | **2** | 5 | 3.951362 | 0.005157 |
| **Base-Seqs** | **1** | 1 | 2.682886 | N/A |
|  | **2** | 1 | 3.925006 | N/A |

The described method allows a flexible, transparent, and clear way to generate novel SSPs. We demonstrated that the method produces similar entropy in sequences as the base set while still preserving accuracy as indicated by Signal-BLAST and MULocDeep. This method could generate peptides with higher yields in synthetic biology or greater uptake of a peptide drug by cells. We believe this method is distinguished from other methods due to its transparent nature. Future work plans include extending the method to create synthetic sequences having multiple biological properties of interest such as targeting of proteins to particular cellular organelles.

## METHODS

### Base sequence data

The base protein sequences used to discover the BBs were obtained from SignalP-5.0 ^11^. Specifically, a total of 2,311 and 7,384 signal and non-signal protein sequences were used, respectively. Each sequence consisted of 70 AAs.

### MLVS model

The method for generating SSPs is based on the previously described MLVS model. The MLVS model represents a protein sequence *S* as a multi-layered collection of ordered *m*-step pairs (*i*,*j*) ∈ ∑, denoted by *P_m|(i,j)_*, *m* = 1,2,…,*k*. The parameter *m* stands for the number of spaces between the elements of the pair, downstream in the flow (left to right) of the sequence, and *k* is the maximum admissible value of *m*. The elements of an ordered pair (*i*,*j*) are referred to as the anchor and tail, respectively, and the location where the anchor element occurs in the sequence is referred to as the anchor location. The elements of the alphabet ∑ are the AAs belonging to a protein sequence. Ordered pairs made up of consecutive elements of the sequence are said to form the family of 1-step pairs, *P_1|(i,j)_*. Allowing multiple spaces between the elements of the ordered pair generates a multitude of *m*-step pairs (families) *P_1_*,*P_2_*,…,*P_m_*,…,*P_k_*, creating a multilayered *k*-clustering *C_k_* made up of sets *P_m|(i,j)_*, *m* = 1,2,…,*k*.

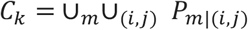

The binding factor between the elements of a particular set *P_m|(i,j)_* is the step size *m*, common for all ordered pairs making up the family. The total number of ordered pairs that can be drawn from the alphabet is |∑|^2^ and the maximum size of C_k_ is reached for k = |S| - 1, where the maximum *m*-step ordered pair (m=k=|S| − 1) spans the entire sequence. Hence, a sequence, S, can be represented as the union of all such ordered pairs at k distinct layers.

In the context of generating SSP sequences, two distinct families of m-step pairs *P_1_*, *P_2_*, …, *P_m_*, …, *P_k_*, are created; one family, *P_1_*, *P_2_*,…, *P_m_*,…, *P_k_*, with respect to the base SP sequences and another family, *P’_1_*,*P’_2_*,…,*P’_m_*,…,*P’_k_*, with respect to the base non-SP sequences. The candidate BBs are identified based on the absolute difference in the frequency of occurrence of individual *m*-step ordered pairs of AAs (*i*,*j*) ∈ ∑ between corresponding sets *P_m_* and *P’_m_*, for *m* = 1,2,…,*k*. The selection of qualified BBs from the candidate BBs and the subsequent process of assembling new SSP sequences follows the steps described above in the main section of the paper.

### Alternative Method for Generating SSPs

An alternative method for generating SSP sequences is to use single AAs as BBs instead of *m*-step ordered pairs of AAs. The twenty canonical AAs (*i.e*., BBs) are sorted in non-decreasing order based on their frequency of occurrence in the given base SP sequences at each location. Assembling a new SSP starts with the selection of a BB at sequence location 1, followed by location 2, and terminating at location (*n* – 1). BBs are selected at each location based on an input parameter, called the *rank-position range*, which represents a range of integer numbers from a specified lower to an upper bound. With respect to the alternative method, the upper bound cannot exceed 20. The rank-position range value selected at a given location *i* determines the BB (*i.e*., AA) at that location.

### Entropy calculations

Entropy and accuracy data were plotted using Prism 8 (for macOS, GraphPad software, San Diego, California USA, www.graphpad.com). To determine if SSP’s were composed of a diverse set of residues, the Shannon entropy formula was applied to the 20 AA alphabet and 20^2^=400 set of bigrams^7^. In each case, the empirical fraction of residues or bigrams was calculated for an input sequence and entered into the function. We elected to use a base 2 logarithm, so entropy units were “nats”.

## Data availability

All data are displayed in the main will be provided in GitHub (https://github.com/) upon acceptance of the manuscript for publication.

## Acknowledgements

We would like to recognize Can Akkoc for his work on the development of the MLVS model and Marlo K. Thompson for her assistance with generating Fig. 1. AP is supported by a National Institutes of Environmental Health Sciences R01 grant #R01ES030084 to AP, and a subcontract of an R35 grant #R35ES031708 to Dr. Joann Sweasy (U. of Arizona).

## Author Contributions

TJ designed the method specifics and implemented the code. RGB assisted with reviewing and updating program source code. GTD performed the entropy calculations. AP assisted with data interpretation and generation of figures. All authors assisted with manuscript preparation and editing.

